# TUNNELING NANOTUBES CONNECT BLOOD-BRAIN BARRIER CELLS: ROLE OF PERICYTES IN BARRIER PRESERVATION DURING ISCHEMIA

**DOI:** 10.1101/2022.03.02.482668

**Authors:** Francesco Pisani, Valentina Castagnola, Laura Simone, Fabrizio Loiacono, Maria Svelto, Fabio Benfenati

## Abstract

Crosstalk mechanisms between pericytes, endothelial cells, and astrocytes preserve integrity and function of the blood-brain-barrier (BBB) under physiological conditions. Long intercellular channels allowing the transfer of small molecules and organelles between distant cells called tunneling nanotubes (TNT) represent a potential substrate for energy and matter exchanges between the tripartite cellular compartments of the BBB. However, the role of TNT across BBB cells under physiological conditions and in the course of BBB dysfunction is unknown.

In this work, we analyzed the TNT’s role in the functional dialogue between human brain endothelial cells, and brain pericytes co-cultured with human astrocytes under normal conditions or after exposure to ischemia/reperfusion, a condition in which BBB breakdown occurs, and pericytes participate in the BBB repair. Using live time-lapse fluorescence microscopy and laser-scanning confocal microscopy, we found that astrocytes form long TNT with pericytes and endothelial cells and receive functional mitochondria from both cell types through this mechanism. The mitochondrial transfer also occurred in multicellular assembloids of human BBB that reproduce the three-dimensional architecture of the BBB. Under conditions of ischemia/reperfusion, TNT formation is upregulated, and astrocytes exposed to oxygen-glucose deprivation were rescued from apoptosis by healthy pericytes through TNT-mediated transfer of functional mitochondria, an effect that was virtually abolished in the presence of TNT-destroying drugs. The results establish a functional role of TNT in the crosstalk between BBB cells and demonstrate that TNT-mediated mitochondrial transfer from pericytes rescues astrocytes from ischemia/reperfusion-induced apoptosis. Our data confirm that the pericytes might play a pivotal role in preserving the structural and functional integrity of BBB under physiological conditions and participate in BBB repair in brain diseases.

## INTRODUCTION

The blood-brain barrier (BBB) functions to protect the brain from the ever-changing peripheral environment and maintain the tightly controlled composition of the brain extracellular fluid necessary for the stability of neuronal activity. Astrocytes, brain endothelium, and pericytes, representing the tripartite structure of the BBB, supervise the blood-brain exchanges and communicate with each other to synergistically produce survival signals required to maintain BBB integrity and central nervous system (CNS) homeostasis.^1, 2^ Paracrine signals between BBB cells, based on diffusible molecules and extracellular vesicles, are known to control BBB functionality,^3, 4, 5, 6, 7^ but the mechanisms governing this intricate intercellular communication network are not fully elucidated.

Alterations of crosstalk mechanisms between BBB cells have been reported under several pathological conditions affecting BBB integrity, such as ischemic stroke.^8, 9^ *In vitro* and *in vivo* experimental evidence shows that ischemia induces alterations of mitochondrial morphology and apoptosis also in astrocytes, contributing to BBB breakdown.^10, 11, 12, 13, 14^ Interestingly, in the post-ischemic phase, pericytes play a pivotal role in the BBB-repairing process.^15^ It is known that pericytes presenting stemness and trans-differentiation potential migrate to the injury sites and contribute to BBB repair.^16^ Furthermore, after CNS injury, pericytes promote the activation of astrocytes participating in damage recovery.^17, 18^

The search for new modalities of communication between BBB cells under physiological conditions and in response to ischemia/reperfusion could contribute to clarifying these mechanisms. Tunneling nanotubes (TNT) are long, F-actin-containing, membranous protrusions connecting distant cells.^19^ TNT allow the intercellular transfer not only of small molecules (such as ions, second messengers, metabolic substrates) but also of macromolecules (proteins and nucleic acids) and organelles, thus generating a cell “support” network that participates in the regulation of tissue functions.^20, 21^ A growing evidence has shown the capability of the TNT-mediated intercellular mitochondria transfer to rescue recipient cells from bioenergetic deficiency and apoptosis, also induced by pathological causes.^22, 23, 24, 25, 26, 27, 28^

Whether TNT play a functional role in the BBB under both physiological and pathological conditions is still unknown. Despite recent data obtained in the mouse retina^29^ and human brain sections^30, 31^ strongly support the possibility that TNT exist at blood-CNS barriers level,^29, 30, 31^ the question of whether BBB cells can communicate with each other through TNT remains open, as well as the question about the possible role of TNT communication between BBB cells during an ischemic insult. Here we demonstrate that BBB cells communicate with each other through TNT and that pericytes and endothelial cells transfer functional mitochondria to normal astrocytes through this mechanism. Similar intercellular trafficking of mitochondria also occurs in an assembloid model of human BBB. Using astrocytes exposed to oxygen-glucose deprivation (OGD) followed by regeneration (OGD/R), an *in vitro* model of ischemia/reperfusion, we demonstrate that TNT transfer mitochondria from healthy pericytes to injured astrocytes to rescue astrocytes from ischemia-induced apoptosis. Similar results were also obtained by directly inducing apoptosis with staurosporine, indicating that this rescue mechanism can also be recruited in the BBB response to other pro-apoptotic stimuli. The identification of TNT between human BBB cells and their activity in promoting intercellular cargo exchanges reveals an important supracellular organization of the BBB that can become essential to preserve its function under pathological conditions.

## RESULTS

### Pericytes transfer functional mitochondria to astrocytes through TNT

We investigated the possibility that astrocytes form heterotypic-TNT junctions with pericytes and that organelles, such as mitochondria, can be exchanged between cell types through this mechanism. To unambiguously identify TNT and distinguish them from other cell-protrusions, three widely accepted criteria were used:^21^ (*i*) TNT are thin and straight membrane protrusions detached from the substrate and connecting two or more cells; (*ii*) they transport and transfer cargoes from a donor to an acceptor cell; (*iii*) they contain F-actin filaments.

All cell cytotypes used were checked for the expression of specific markers prior experiments (**Suppl. Figure 1**). Intercellular trafficking of mitochondria was analyzed using astrocytes as acceptor cells and pericytes as donor cells. To this aim, primary astrocytes were stained with CellMask Orange (CM) and co-cultured, at a low confluence, with primary pericytes previously stained with ΔΨ-dependent MitoTracker Deep Red (MT). Co-cultures were analyzed for mitochondrial trafficking from 1 to 24 h after seeding by live time-lapse fluorescence microscopy and laser-scanning confocal microscopy (**Figure 1A**). We observed that pericytes transferred mitochondria to astrocytes by direct cell-to-cell contact mainly in the first hours after seeding (1-4 h; **Figure 1A**, panel A1; **Suppl. Movie 1**). Starting from 1 h after seeding (**Figure 1A**, panel A2; **Suppl. Movie 2**) and up to 24 h (**Figure 1A**, panels A3, A4; **Suppl. Movies 3-7**; **Figure 1B**), we mostly observed the formation of long intercellular plasma membrane bridges that appeared detached from the glass surface, through which labeled mitochondria were actively transported from pericytes to astrocytes. These structures were most likely TNT, meeting two out of the three identification criteria reported above.

**Figure 1.**
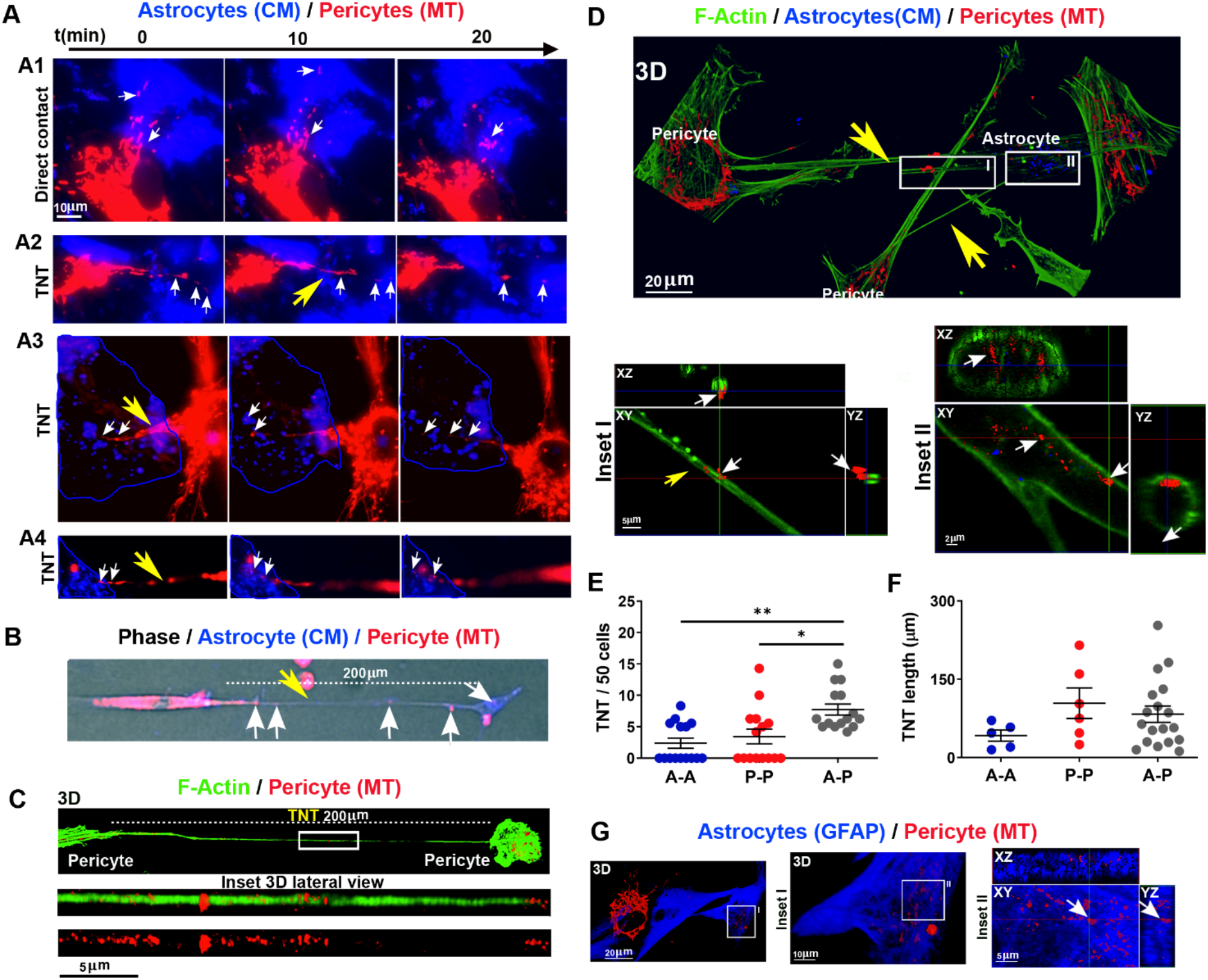
Pericytes transfer functional mitochondria to astrocytes via TNT. Functional mitochondria of pericytes were stained with ΔΨ-dependent MitoTracker Deep Red (MT; red), while the plasma membrane of astrocytes was stained with CellMask Orange (CM; blue). Cells were co-cultured and analyzed for intercellular trafficking of mitochondria by time-lapse and confocal fluorescence microscopy. **A.** Live time-lapse analysis of mitochondria trafficking. Immediately after seeding, MT-labeled mitochondria (red) from pericytes are actively transferred to astrocytes at direct contacts (panel A1; Suppl. Movie 1). After 1 h (panel A2; Suppl. Movie 2) and up to 24 h (panels A3, A4; Suppl. Movie 3-7), long plasma membrane intercellular bridges, a.k.a. TNT, are formed through which mitochondria are transferred from pericytes to astrocytes. After a few hours, the plasma membrane marker of astrocytes undergoes internalization in the form of intracellular vesicles (blue) due to the ongoing membrane turnover. Yellow arrows, TNT; white arrows, mitochondria. Representative images from N = 7 independent preparations, with >10 sampled areas per experiment. **B.** Widefield imaging of a long pericyte-astrocyte TNT. TNT (200 μm in length; yellow arrow) between a pericyte (red, left) and an astrocyte (blue, right) acquired by live widefield fluorescence microscopy. Note the presence of pericyte mitochondria (white arrows) inside the TNT. **C.** Confocal imaging of pericyte-pericyte TNT. After 24 h of co-culture, cells were fixed and stained for F-actin using fluorescent phalloidin (green). 3D reconstruction of a long (> 200 μm) pericyte-pericyte TNT in which many mitochondria are visible. **D.** 3D reconstruction of a long pericyte-astrocyte TNT. The long TNT stained for F-actin using fluorescent phalloidin (green) is detached from the glass surface and connects to an astrocyte along which mitochondria from the pericyte are visible (inset I; red). The lower inset (inset II) shows an intracellular XY plane of the CM-labeled astrocyte (blue intracellular vesicles) in which many mitochondria of pericyte origin are visible (red). Representative images from N=3 independent experiments with > 10 sampled areas per experiment. **E.** Quantification of homotypic-TNT (A-A or P-P) and heterotypic-TNT (A-P) number after 24 h of co-culture. A single dot represents the TNT number/field of 50 cells. N=3 independent experiments with > 10 sampled areas per experiment. **F.** Quantification of homotypic-TNT and heterotypic-TNT length after 24 h of co-culture. N=3 independent experiments with > 10 sampled areas per experiment. **G.** GFAP immunofluorescence analysis. 3D reconstruction, XY plane, and Z-projections by confocal microscopy show pericyte mitochondria inside GFAP-positive astrocytes. Representative images from N=3 independent experiments with > 10 sampled areas per experiment. In E,F: N = 3 independent experiments with > 10 sampled areas per experiment. *p<0.05, **p<0.005; Kruskal-Wallis/Dunn’s tests.

We next fixed the co-cultures and performed confocal microscopy analysis to evaluate whether these putative TNT were F-actin positive, the exact position of these structures with respect to the glass surface, and the localization of pericyte-derived mitochondria inside astrocytes. Indeed, pericyte-derived mitochondria were observed inside straight F-actin-positive structures that, running in suspension over the glass surface, were connecting pericyte to pericyte (homotypic-TNT, **Figure 1C**) or pericyte to astrocytes (heterotypic-TNT, **Figure 1D**, inset I). Furthermore, several pericyte-derived mitochondria were found inside astrocytes, together with CM-positive intracellular vesicles resulting from the progressive internalization of stained plasma membrane patches (**Figure 1D**, inset II). Quantitative analysis of homotypic- and heterotypic-TNT revealed that the frequency of heterotypic-TNT was significantly higher with respect to homotypic-TNT (**Figure 1E**). High heterogeneity was observed for both pericyte-pericyte and heterotypic TNT lengths (range: 15-253 μm; **Figure 1F**).

To further confirm that the transfer of functional mitochondria from donor pericytes was directed to astrocytes, we immunolabeled samples for the astrocyte-specific marker glial fibrillay acid protein (GFAP). Confocal microscopy analysis confirmed that pericytes’ mitochondria were transferred inside GFAP-positive astrocytes (**Figure 1G**).

### Endothelial cells transfer functional mitochondria to astrocytes through TNT

A similar picture involving the formation of heterotypic TNT and intercellular trafficking of mitochondria was observed when using endothelial cells as donors and astrocytes as acceptors (**Figure 2**; **Suppl. Movies 8-9**). CM-stained astrocytes were co-cultured with MT-labeled endothelial cells and analyzed for mitochondrial trafficking by live time-lapse fluorescence microscopy and laserscanning confocal microscopy, as described above (**Figure 2A**). We observed that, after a transient phase of direct cell-to-cell mitochondria transfer (1 h of co-culture; **Figure 2A**, panel A1; **Suppl. Movie 8**), endothelial cells transferred mitochondria to astrocytes through the formation of long intercellular TNT up to 24 h after seeding (**Figure 2A**, panels A2-A4; **Suppl. Movie 9**). Next, using confocal microscopy analysis of fixed co-cultures, we confirmed that the putative TNT were positive for F-actin and detached from the glass surface and that mitochondria of endothelial origin were present inside astrocytes (**Figure 2B,C**). Confocal analysis with 3D reconstruction revealed that endothelial mitochondria were traveling along heterotypic TNT to reach GFAP-positive acceptor astrocytes (**Figure 2D**). Quantitative analysis of homotypic- and heterotypic-TNT frequency revealed that, opposite to the astrocyte/pericyte co-culture reported above, the frequency of heterotypic-TNT was similar to those of homotypic-TNT (**Figure 2E**), with heterogeneous lengths of the TNT (range: 16-207 μm; **Figure 2F**).

**Figure 2.**
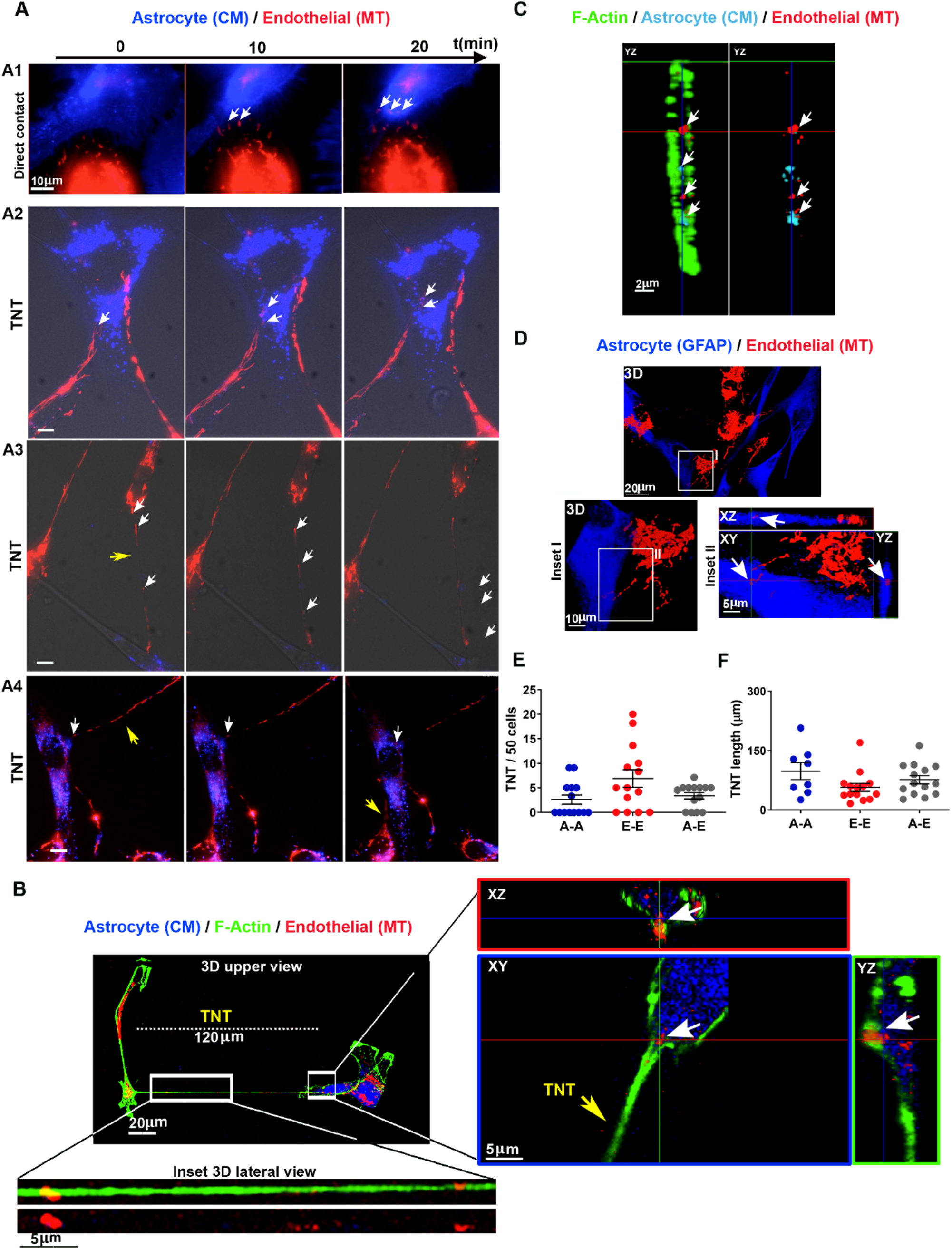
Endothelial cells transfer functional mitochondria to astrocytes via TNT. Functional mitochondria of endothelial cells were stained with MT (red), while astrocyte membranes were labeled with CM (blue). Live-stained cells were co-cultured and analyzed for intercellular mitochondria transport by time-lapse and confocal fluorescence microscopy. **A.** Live-cell time-lapse analysis of mitochondrial trafficking. One hour after seeding, red-mitochondria (white arrows) from endothelial cells were actively transferred to astrocytes at cell-to-cell contacts (panel A1, Suppl. Movie #8). After a few hours and up to 24-48 h, long intercellular TNT connected astrocytes with endothelial cells and transferred mitochondria (white arrows) from endothelial cells to astrocytes (A2-A4; Suppl. Movie #9). Representative frames from N = 7 independent preparations, with >10 sampled areas per experiment. **B.** Confocal imaging of a long endothelial cell-astrocyte TNT. 3D reconstruction of XY confocal planes of a 120 μm-long TNT connecting an astrocyte (GFAP, blue) to an endothelial cell. The magnified inset reported at the bottom, shows the 3D reconstruction of endothelial mitochondria (red) traveling along the TNT (actin, green). The magnified inset reported on the right shows a single XY plane and XZ/YZ projections of a TNT attached to the astrocyte in which endothelial mitochondria are localized. **C.** Confocal microscopy analysis of F-actin (green), astrocyte plasma membrane and internalized vesicles (cyano), and endothelial mitochondria (red). Note the presence of endothelial-derived mitochondria inside the astrocyte. Representative frames from N = 3 independent preparations, with > 10 sampled areas per experiment. **D.** GFAP immunofluorescence analysis. 3D reconstruction, XY plane and Z-projections show the presence of endothelial mitochondria inside a GFAP-positive astrocyte. Representative images from N = 3 independent preparations, with > 10 sampled areas per experiment. **E.** Quantification of homotypic-TNT (A-A or E-E) and heterotypic-TNT (A-E) number after 24 h of co-culture. A single dot represents the TNT number/field of 50 cells. **F.** Quantification of homotypic and heterotypic TNT length after 24 h of co-culture. In E,F: N = 3 independent experiments with > 10 sampled areas per experiment. Kruskal-Wallis/Dunn’s tests.

### Pericytes and endothelial cells transfer mitochondria to astrocytes in a 3D assembloid model of human BBB

To test the hypothesis that mitochondria deriving from either pericytes or endothelial cells are transferred to astrocytes in a more physiological model of human BBB, we generated human BBB organoids/multicellular assembloids, as previously reported.^32^ BBB assembloids were checked for barrier protein expression and BBB integrity (**Suppl. Figure 2**). Brain endothelial cells, pericytes, and astrocytes spontaneously assembled in core-shell spheroids with a core entirely made of astrocytes and an external shield made of a continuous layer endothelial cells, sealing the assembloid and separating it from the external environment. The penetration of these spheroids by two prototypic molecules known not to permeate the BBB (4-kDa fluorescent dextran – paracellular transport) or be efficiently transported through it (fluorescent transferrin – transcellular transport) showed that the BBB assembloids were a reliable BBB model (**Suppl. Figure 2**).

To assess the intercellular mitochondria transfer by confocal microscopy, functional BBB assembloids were generated starting from CM-stained astrocytes and either pericytes or endothelial cells whose mitochondria were stained with ΔΨ-dependent MT. After 48 h, assembloids were fixed and imaged at low magnification by confocal microscopy. The strongest MT signal was found in the peripheral region of the assembloid, particularly when mitochondria from endothelial cells were primarily stained, while CM-stained astrocytes were found in the central region (**Figure 3A**, panel A1; **Figure 3B**, B1). Interestingly, the 3D reconstruction of confocal planes acquired at high magnification showed the presence of MT signal in the central region, suggesting that mitochondria of pericytes (**Figure 3**, panels A2-A3) or endothelial cells (**Figure 3B**, panels B2-B3) reached the astrocytic core of the BBB assembloid. To confirm the actual transfer of mitochondria to the cytoplasm of astrocytes, the whole assembloids were fixed and analyzed by immunofluorescence using anti-GFAP antibodies and focusing on the confocal planes corresponding to the assembloid core (50 μm depth). Mitochondria of endothelial and pericytic origin were found inside GFAP-positive astrocytes when the assembloids were generated using MT-labeled endothelial cells and pericytes, respectively (**Figure 3C,D**).

**Figure 3.**
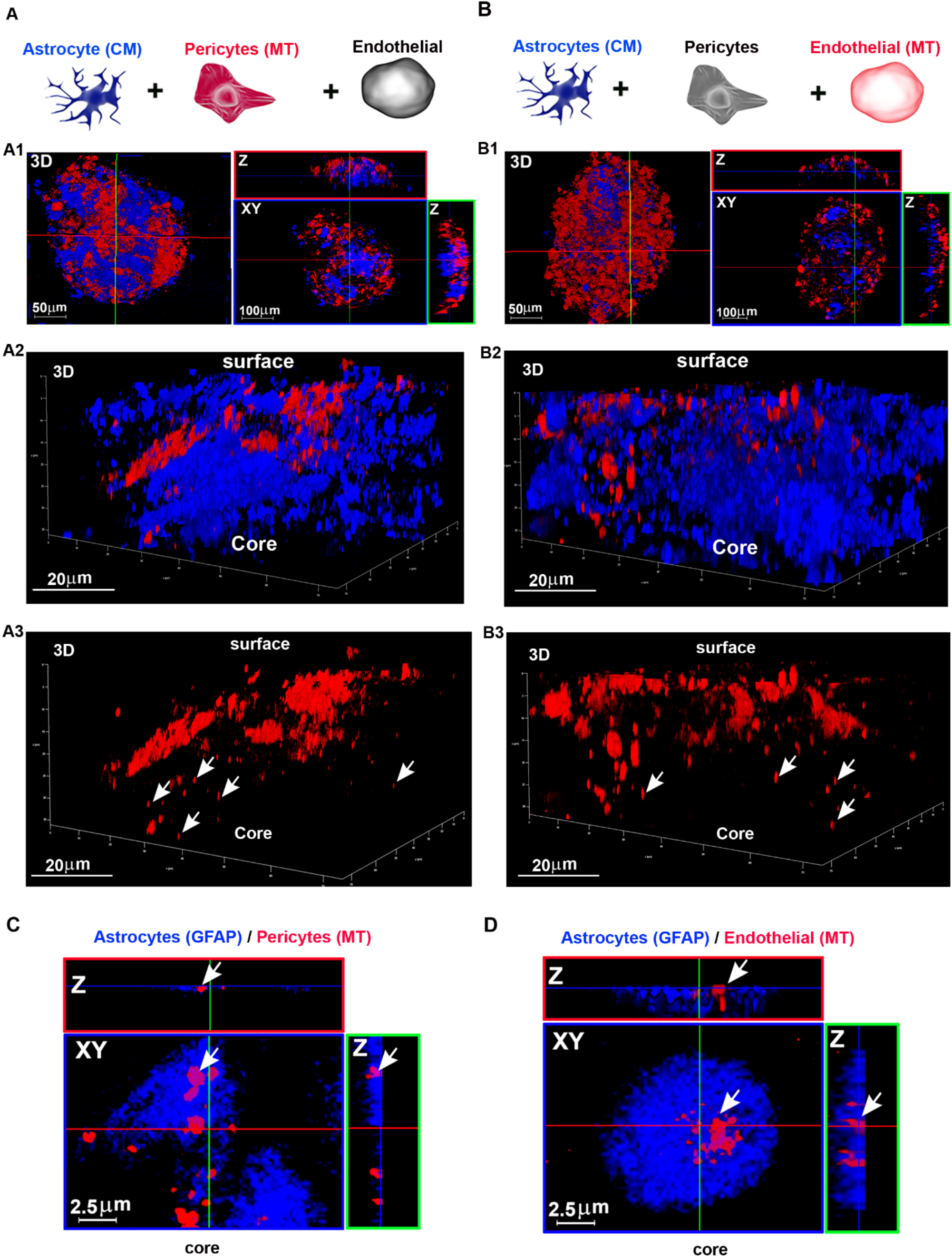
Endothelial cells and pericytes transfer functional mitochondria to astrocytes in a 3D assembloids model of human BBB. **A,B.** Functional mitochondria from either pericytes or brain endothelial cells were stained with MT (red), while astrocyte membranes were labeled with CM (blue). To generate BBB assembloids, labeled astrocytes were cultured with either labeled pericytes and unlabeled endothelial cells (**A**) or labeled endothelial cells and unlabeled pericytes (**B**). After 2 days in culture, assembloids were fixed and analyzed by confocal microscopy for MT localization. 3D reconstructions and Z-projections acquired at low magnification show either pericytes (A1, red) or endothelial cells (B1, red) localized in the assembloid shell, with astrocytes localized in the assembloid core (A1, B1, blue). In both cases, high magnification analysis reveal MT-stained mitochondria (red) in the assembloid core (A2/A3; B2/B3). Representative images from N = 3 independent preparations with N ≥ 3 assembloids per experiment. **C,D.** Immunofluorescence and high magnification confocal analysis of the assembloid core using anti-GFAP antibodies. XY planes and Z projections of the assembloid core show MT-stained mitochondria (red) from either pericyte (**C**) or endothelial cells (**D**) inside GFAP-positive astrocytes. Representative images from N = 2 independent experiments in which N ≥ 6 assembloids per experiment were analyzed.

### Oxygen-glucose deprivation/reoxygenation boosts TNT formation between astrocytes and healthy pericytes

TNT are frequently upregulated in response to cellular stress to restore damaged functions in receiving cells.^33^ Thus, we tested whether astrocytes exposed to hypoxia/reoxygenation stress, an *in vitro* model of ischemia/reperfusion^34^ known to induce mitochondrial damage in astrocytes,^35^ were able to induce the formation of an increased number of TNT with healthy pericytes and whether the F-actin cytoskeleton plays a role in this response and in the subsequent mitochondria transfer.

Astrocytes were stained with CM and exposed to 2% O_2_ in glucose- and serum-free medium for 24h (oxygen-glucose deprivation, OGD). After OGD, astrocytes were washed and regenerated (OGD/R) in complete medium with a constant number of healthy MT-labeled pericytes in the presence or absence of the F-actin-depolymerizing agent cytochalasin D (CytoD). After 24 h of OGD/R, the coculture was fixed, stained for F-actin with phalloidin-AlexaFluor488, and analyzed by confocal microscopy to quantify the number of astrocyte-pericyte TNT and the presence of pericyte-derived mitochondria inside astrocytes (**Figure 4A**).

**Figure 4.**
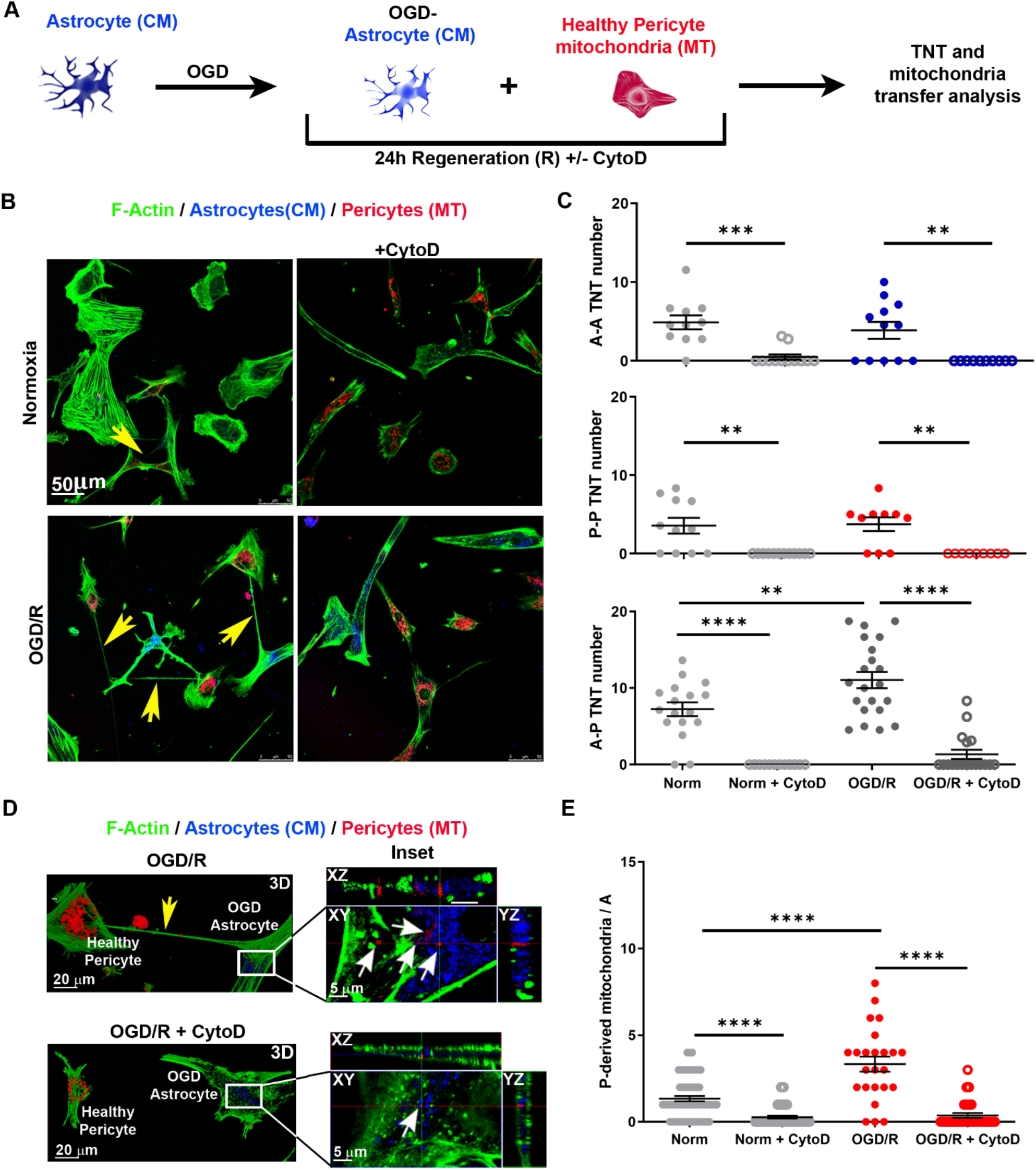
Oxygen-glucose deprivation/reoxygenation boost TNT formation between astrocytes and healthy pericytes in a cytoskeleton-dependent manner. **A.** Schematic representation of the oxygen-glucose deprivation/reoxygenation (OGD/R) paradigm. Astrocytes, stained with CM and exposed to 2% O_2_ for 24 h in glucose- and serum-free medium (OGD), were co-cultured for 24 h (regeneration phase) with a constant number of healthy MT-stained pericytes in the presence or absence of Cytochalasin D (CytoD; 200 nM). Cells were fixed and analyzed by confocal microscopy for TNT structure and number, phalloidin-labeled F-actin, and mitochondria transfer. **B.** OGD-astrocytes trigger heterotypic-TNT formation with healthy pericytes. Astrocyte-to-pericyte TNT (yellow arrows) are indicated in normoxia and in OGD/R. Note that OGD/R trigger TNT while CytoD completely prevents TNT formation. Images are representative of N = 4 independent experiments. **C.** Quantitative evaluation of the astrocyte-astrocyte (A-A), pericytepericyte (P-P) and heterotypic astrocyte-pericyte (A-P) number of TNT after 24 h of co-culture. OGD/R strongly upregulates heterotypic-TNT formation, while CytoD completely prevents TNT formation in control and OGD/R conditions. A single dot represents the TNT number/field of 50 cells. N = 3; ** p<0.005 *** p<0.001, ****p<0.0001; two-way ANOVA/Tukey’s tests. **D.** 3D confocal reconstruction at higher magnification of the experiments shown in B. XY plane and Z-projections show a representative [OGD/R astrocytes + pericytes] co-culture incubated in the absence (*up*) or presence of CytoD (*bottom*). TNT between OGD/R astrocyte and pericyte and pericyte-derived mitochondria are indicated by yellow and white arrows, respectively. Note that CytoD treatment prevents the formation of TNT and strongly reduces mitochondria transfer from pericytes to astrocytes. Images are representative of N = 4 independent experiments. **E.** Quantification of pericyte-to-astrocyte mitochondrial transfer. N = 3; ****p<0.0001; two-way ANOVA/Tukey’s tests.

TNT between OGD/R-astrocytes and pericytes were significantly more numerous than those observed under normoxic conditions, indicating that astrocyte OGD/R triggers higher TNT formation with pericytes. CytoD treatment completely prevented TNT formation in both control and OGD/R co-culture, showing the pivotal role of F-actin cytoskeleton in this mechanism (**Figure 4B,C**).

High-resolution confocal microscopy with 3D reconstruction, followed by quantitative analysis of mitochondrial transfer, indicated that OGD/R triggers the translocation of mitochondria from pericytes to astrocytes, while the lack of TNT due to CytoD treatment was associated with a reduced number of pericyte-derived mitochondria in both control and OGD/R-astrocytes (**Figure 4D,E**). Overall, the data indicate that astrocytes stressed by OGD/R trigger TNT formation with healthy pericytes in a cytoskeleton-dependent manner that act as a structural substrate for the intercellular mitochondrial transfer.

### TNT-mediated transfer of mitochondria from pericytes to astrocytes rescues OGD/R-induced astrocyte apoptosis

After the demonstration that OGD/R enhances the formation of astrocytes-pericytes TNT and the transfer of pericytes’ mitochondria in stressed astrocytes, we investigated whether this phenomenon could lead to the rescue of astrocytes from OGD/R apoptosis, as previously observed for other cell types.^22, 23, 24, 25, 26, 27^ To this aim, CM-labeled astrocytes were exposed to OGD, washed and reoxygenated in normal complete medium for 24 h (OGD/R) with a constant number of healthy pericytes in the presence or absence of CytoD. The co-culture was then analyzed for the extent of mitochondria trafficking and astrocytes apoptosis.

Live time-lapse analysis confirmed the movement of mitochondria from pericytes to OGD-astrocytes that was dependent on microfilament integrity (**Figure 5A**, panel 1, insets I-III; **Suppl. Movie 10**) and was not observed in the presence of CytoD (**Figure 5A**, panel 3, insets I-III; **Suppl. Movie 12**). Mitochondrial transfer also occurred between healthy pericytes and OGD-astrocytes with overt morphological alterations (**Figure 5A**, panel 2, insets I-III; **Suppl. Movie 11**). In both cases, CytoD abolished TNT formation and mitochondrial transfer (**Figure 5A**, panel 4, insets I-III; **Suppl. Movie 13**).

**Figure 5.**
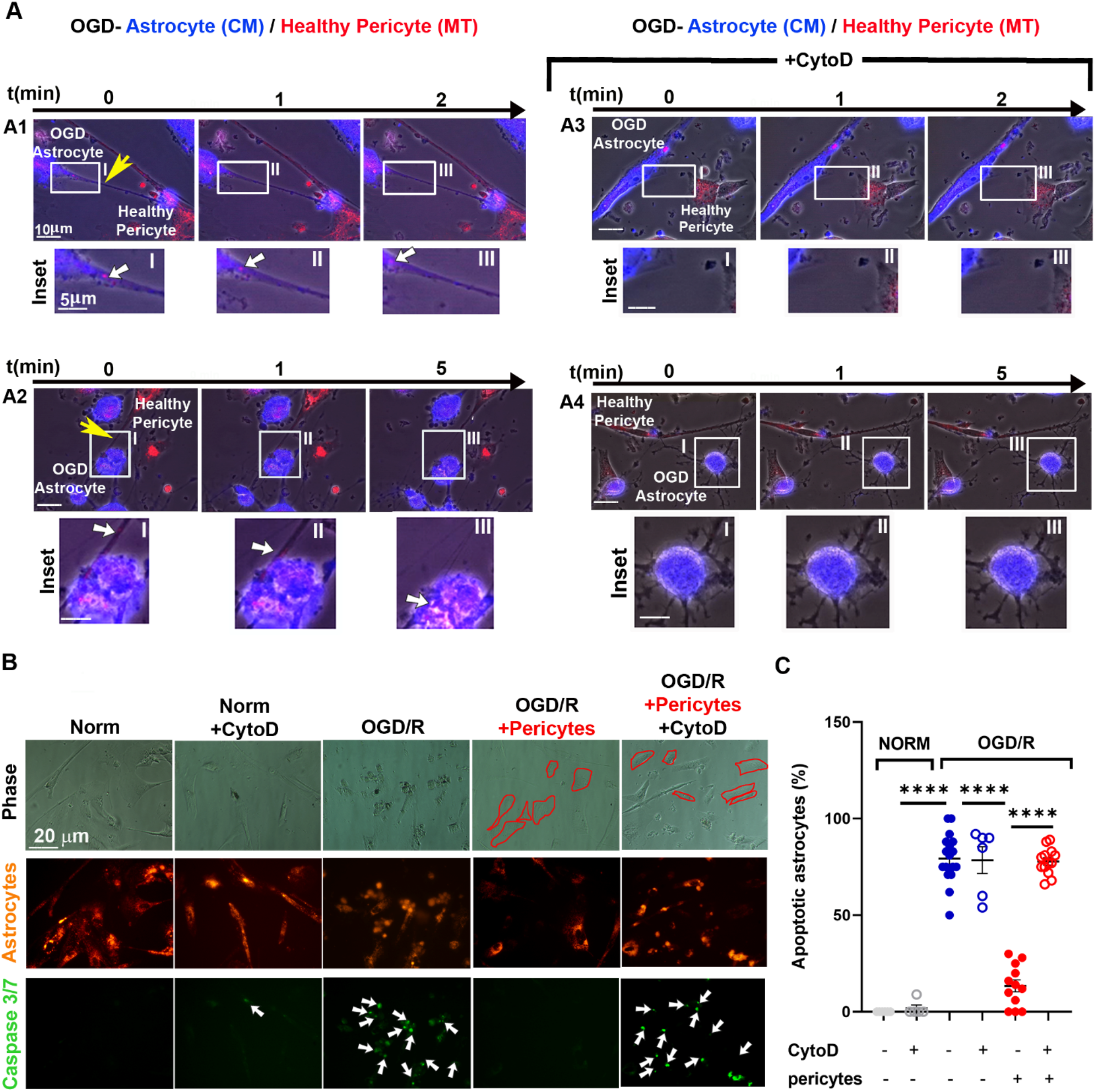
TNT-mediated mitochondrial transfer from pericytes rescues OGD-induced apoptosis of astrocytes. Astrocytes were stained with CM and exposed to 2% O_2_ for 24 h in glucose- and serum-free medium, washed and refreshed in normal complete medium with a constant number of healthy MT-stained pericytes (OGD/R), in the presence or absence of CytoD (200 nM), as described in Figure 4A. Twenty-four hours later, the co-culture was analyzed for mitochondria trafficking and astrocyte apoptosis by time-lapse and confocal fluorescence microscopy. **A.** Time-lapse fluorescence microscopy analysis of mitochondria trafficking from healthy pericytes to OGD-astrocytes regenerated for 24 h (OGD/R) in the absence or presence of CytoD. TNT and actively transported mitochondria from healthy pericytes to OGD-astrocytes are indicated by yellow and white arrows, respectively (A1 and Suppl. Movie 10; A2 and Suppl. Movie 11 for OGD-astrocytes with overt morphological alterations). CytoD abolished TNT formation and strongly reduced the intercellular mitochondria trafficking to OGD-astrocytes (A3; Suppl. Movie 12) also when astrocytes are morphologically altered (A4; Suppl. Movie 13). Representative images from N = 4 independent experiments in which N ≥ 10 areas per experiment were analyzed. **B.** Apoptosis of OGD-astrocyte regenerated for 24 h (OGD/R) with healthy pericytes in the presence or absence of CytoD. Astrocytes (orange) that were positive for CellEvent Caspase 3/7 (green nuclei, white arrows) were counted under the experimental conditions reported in the images. **C.** Analysis of OGD-astrocyte apoptosis. The percentage of Caspase 3/7-positive astrocytes on the total astrocyte population is shown. Pericytes rescued astrocytes from OGD-induced apoptosis only in the presence of an intact F-actin cytoskeleton. N = 3 independent experiments. ****p<0.0001; Student’s t-test (normoxia); two-way ANOVA/Tukey’s tests (OGD/R).

The same co-culture was analyzed for the percentage of apoptotic astrocytes by adding CellEvent caspase 3/7 green to the medium and evaluating caspase 3/7-positive astrocytes by fluorescence microscopy. This analysis showed that OGD/R induces significant apoptosis in astrocytes and that the presence of pericytes during post-OGD reoxygenation strongly reduced the percentage of apoptotic astrocytes (**Figure 5B,C**). The treatment with CytoD completely prevented the rescue of astrocytes from apoptosis indicating a causal link between TNT formation, mitochondria transfer, and rescue from apoptosis (**Figure 5B,C**).

### Transfer of mitochondria from pericytes through TNT rescues astrocytes from induced apoptosis induced by staurosporine treatment

To investigate whether the formation of heterotypic TNT and mitochondrial transfer is a general BBB response to cellular stress astrocytes were treated with staurosporine (STS, 1 μM for 3 h), another cell stressor affecting the functionality of mitochondria^36^, and regenerated in an STS-free medium with healthy pericytes (**Figure 6A**). As observed for the OGD/R protocol, the STS/R treatment significantly increased the number of pericyte-astrocyte TNT (**Figure 6B,C**), as well as the presence of pericyte mitochondria inside STS/R-astrocytes (**Figure 6D,E**). Similar to OGD/R, also in this case, the actin-depolymerizing agent CytoD fully inhibited TNT formation and the related mitochondrial exchange (**Figure 6B-E**). Interestingly, also the treatment with the microtubule-depolymerizing drug vincristine (VCR) significantly reduced the formation of pericyte-astrocyte TNT (**Suppl. Figure 3**). Live time-lapse analysis showed that pericytes transferred mitochondria mainly through TNT to STS-astrocytes (**Figure 7A**, panel 1, insets I-III; **Suppl. Movies 14,15**) and that this transfer was prevented by VCR (**Figure 7A**, panel 2, insets I-III; **Suppl. Movie 16).** The analysis of the extent of apoptosis by measuring the caspase 3/7 activity showed that pericytes rescue apoptotic STS-astrocytes and that VCR strongly reduces this rescue (**Figure 7B,C**). Similar results were obtained also analyzing necrotic astrocytes detached from glass surface through flow cytometry (**Suppl. Figure 4**). Overall, the data indicate that pericytes rescue astrocytes from STS-induced apoptosis through TNT-mediated mitochondrial transfer that depends on the integrity of the cytoskeleton.

**Figure 6.**
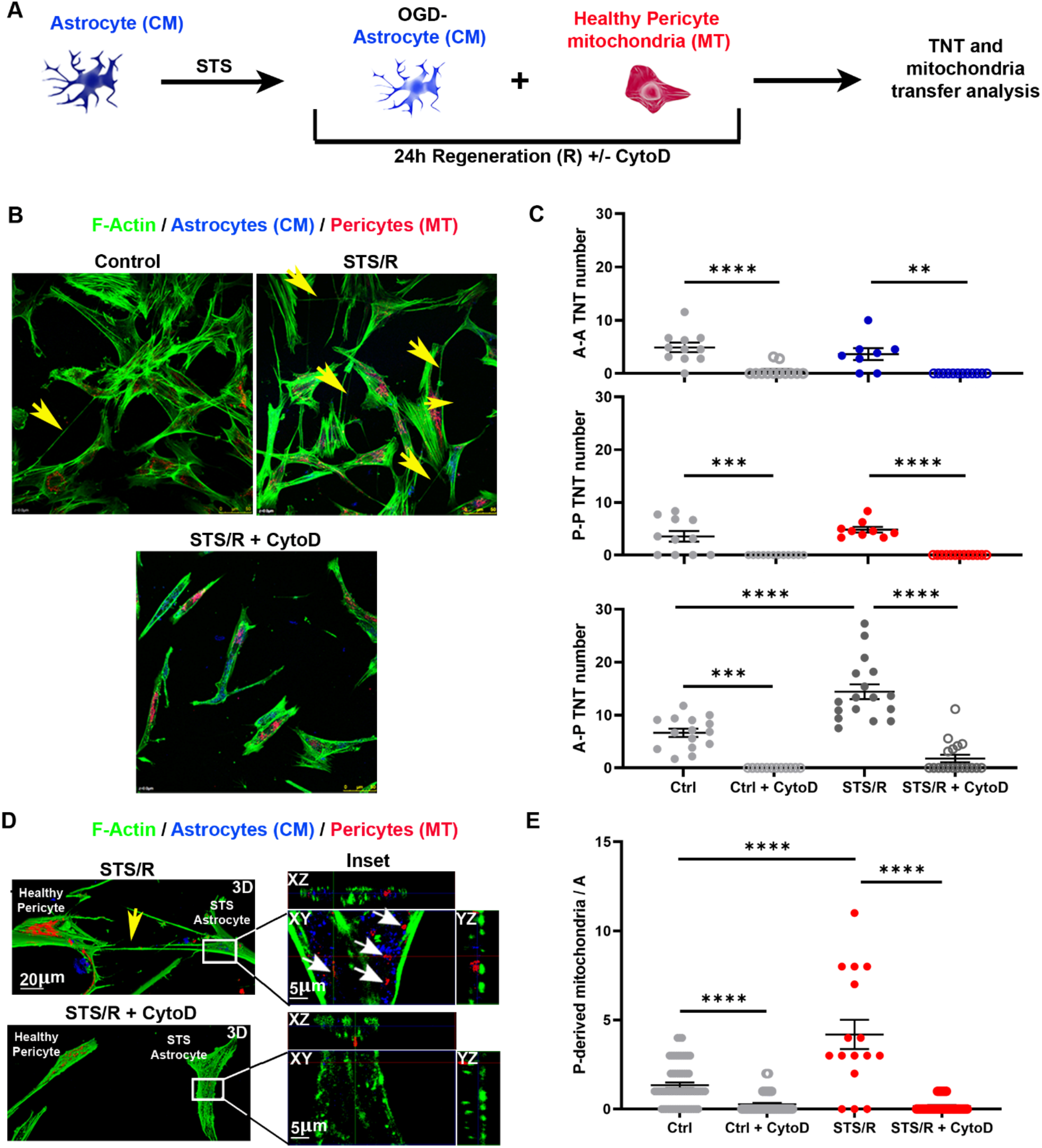
Staurosporin treatment of astrocytes triggers TNT formation with healthy pericytes in a cytoskeleton-dependent manner. **A.** CM-labeled astrocytes (blue) were treated with STS (1 μM for 3 h), washed, and co-cultured with pericyte that had been previously stained with MT (red) (STS/R) with or without CytoD (STS/R+CytoD). After 24 h, cells were fixed, stained for the F-actin-network with phallodin (green), and analyzed by confocal microscopy for the TNT number and mitochondria transfer. **B.** STS-astrocytes trigger TNT heterotypic-TNT formation with healthy pericytes. Astrocyte-to-pericyte TNT (yellow arrows) are indicated in control and in STS/R. Note that STS/R trigger TNT while CytoD completely prevents TNT formation. Homotypic (A-A and P-P) and heterotypic (A-P) TNT were counted between control or STS-astrocytes and pericytes (STS/R) in the absence or in the presence of CytoD (STS/R+CytoD). **C.** Quantitative evaluation of the astrocyte-astrocyte (A-A), pericyte-pericyte (P-P) and heterotypic astrocyte-pericyte (A-P) number of TNT after 24 h of co-culture. STS-astrocytes strongly upregulate heterotypic-TNT formation, an effect that is strongly inhibited by CytoD treatment. **D.** 3D confocal reconstruction at higher magnification of the experiments shown in D. XY plane and relative Z-projections show a representative [STS/R astrocytes + pericytes] co-culture incubated in the absence (*up*) or presence (*bottom*) of CytoD. TNT between STS/R astrocytes and pericytes are indicated by the yellow arrows; pericyte-derived mitochondria are indicated by the white arrows. Note that CytoD treatment prevents the formation of TNT and strongly reduces mitochondria transfer from pericytes to astrocytes. Images are representative of N = 4 independent experiments. **E.** Quantification of pericyte-to-astrocytes mitochondria transfer. N = 3; ** p<0.005 *** p<0.001, ****p<0.0001, twoway ANOVA/Tukey’s tests.

**Figure 7.**
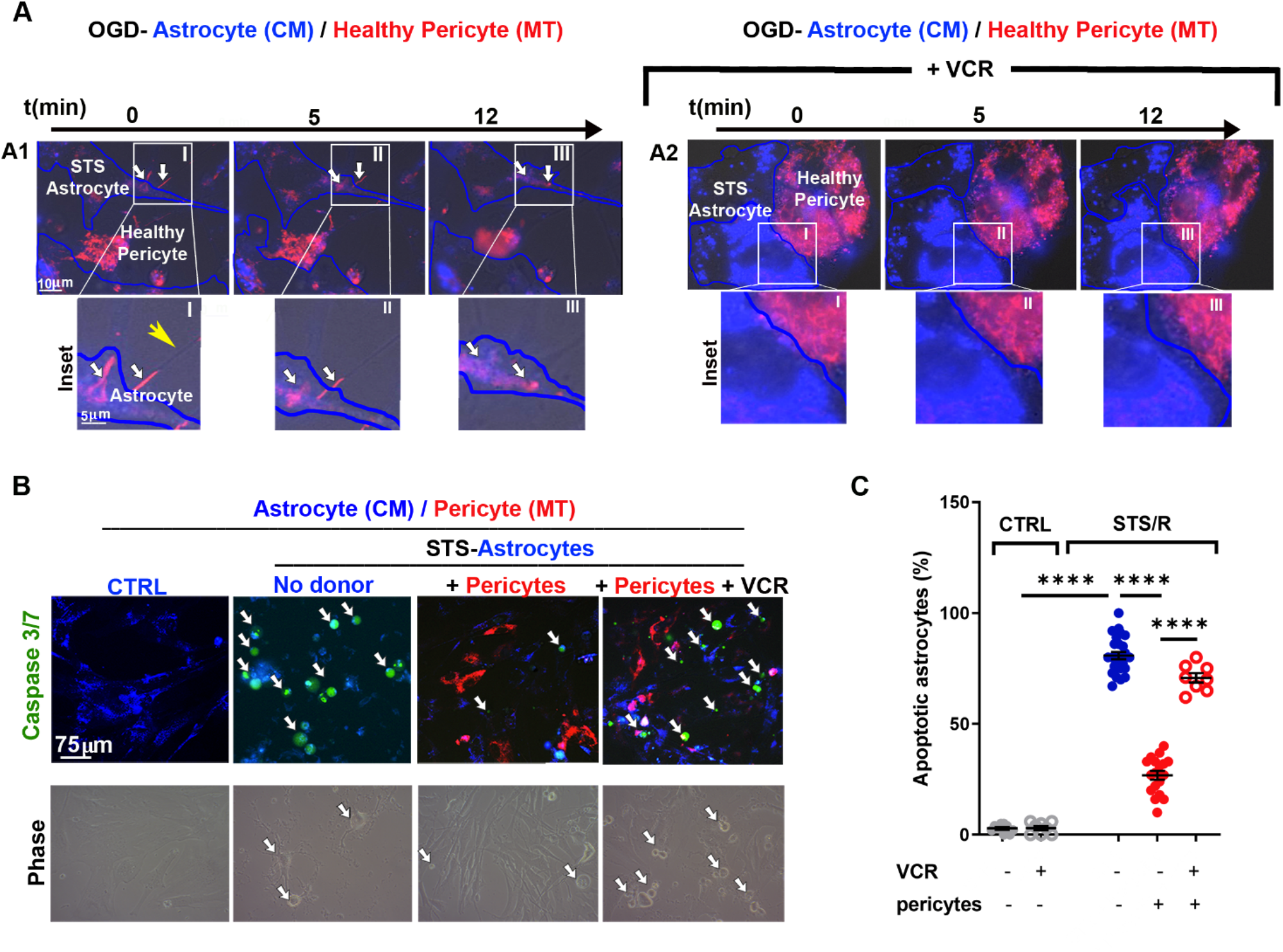
Pericytes rescue STS-induced astrocyte apoptosis by TNT-mediated transfer of mitochondria. CM-stained astrocytes were incubated with STS for 3 h. After the treatment, cells were washed and co-cultured for 24 h with pericytes (+Peri) as described in Figure 6A, in the presence or absence of the tubulin-depolymerizing drug vincristine (VCR; 10 nM). The co-culture was analyzed for mitochondria trafficking and apoptosis by time-lapse and confocal fluorescence microscopy. **A.** Time-lapse analysis of mitochondrial trafficking from healthy pericytes to STS-treated astrocytes. White arrows indicate mitochondria actively transported from healthy pericytes to STS-astrocytes through TNT (yellow arrow) (panel A1, insets I-III; Suppl. Movie 14-15). VCR treatment prevents TNT formation and strongly reduces intercellular mitochondrial trafficking (panel A2, insets I-III; Suppl. Movie 16). Representative images from N = 17 independent experiments with N ≥ 10 areas per experiment analyzed. **B.** Apoptosis of STS-treated astrocytes regenerated for 24 h with healthy pericytes in the presence or absence of VCR. Astrocytes (blue) that were positive for CellEvent Caspase 3/7 (green nuclei, white arrows) were counted under the experimental conditions reported in the images. **C.** Analysis of STS-astrocyte apoptosis. The percentage of Caspase 3/7-positive on the total astrocyte population is shown. Pericytes strongly rescue astrocytes from STS-induced apoptosis, an effect that is significantly decrease by VCR. N = 4 independent experiments. ****p<0.0001; Student’s t-test (control); two-way ANOVA/Tukey’s tests (STS/R).

## DISCUSSION

Intercellular crosstalk is fundamental in supracellular organizations, such as the BBB. Here, we describe a new modality on intercellular communication based on functional heterotypic-TNT, through which pericytes and endothelial cells communicate with astrocytes by exchanging energy and matter. We characterized the structure and function of these TNT that, in addition to informational and metabolic exchanges, were found to rescue astrocytes from apoptosis through the intercellular transfer of functional mitochondria from healthy pericytes to ischemic/apoptotic astrocytes. Furthermore, we provide evidence that mitochondria transfer also occurs in 3D multicellular assembloids of human BBB.

It is well known that mitochondria are pivotal organelles, not only for oxidative metabolism, but also for cell signaling, calcium homeostasis, cell differentiation, proliferation, death, and reprogramming of differentiated cells.^37, 38, 39, 40^ Consequently, the transcellular transfer of functional mitochondria potentially modifies these functions in recipient cells. In particular, it has been shown that the intercellular mitochondria transfer participates in tissue homeostasis under physiological conditions and contributes to the repair of injured tissues.^41^ Mitochondria have been described to migrate from one cell to another through distinct mechanisms. Among those, extracellular vesicles, gap junctions, exocytosis and endocytosis of naked mitochondria, cytoplasmic fusion, and TNT have been reported.^42, 43, 44^ In our co-culture model, heterotypic-TNT were found to be the primary mechanism by which functional mitochondria were transferred between either endothelial cells or pericyte to distant astrocytes.

We investigated whether this phenomenon represents a call-for-help mechanism aimed at rescuing astrocytes from apoptosis, e.g., after ischemia/reperfusion, as previously shown for other types of recipient cells.^22, 23, 24, 25, 26, 27^ Although both endothelial cells and pericytes were very efficient in transferring functional mitochondria to astrocytes, we focused our attention on pericytes due to their role in central nervous system regeneration after ischemic stroke.^6, 15, 30, 45, 46, 47^ Indeed, pericytes have been identified as a new therapeutic target in ischemic stroke^45^ and have been proposed as a valuable tool in BBB-regenerative medicine approaches.^48, 49^ In this respect, the same pericyte cell line (hBVP) used in our work was found to ameliorate BBB integrity, prevent neuronal apoptosis, and improve the neurological function when transplanted in a mouse model of middle cerebral artery occlusion with BBB breakdown.^15^ Our data show that OGD/R-injured astrocytes strongly trigger TNT formation with healthy pericytes, suggesting that the biogenesis of TNT is a call-for-help mechanism aimed at rescuing cells from apoptosis. Based on our data, it is tempting to speculate that during the post-ischemic BBB repair process, pericytes migrate to the injured site and rescue astrocytes from ischemia through TNT-mediated mitochondria transfer. This could contribute both to astrocyte survival and post-ischemic BBB repair. The restorative role of pericytes is consistent with their multipotential stem cell activity and similarity with mesenchymal stem cells (MSCs).^50, 51, 52^ Interestingly, MSCs rescue apoptotic cells through the TNT-mediated mitochondria transfer, as shown here for pericytes.^22, 23, 24, 25, 26^

The cytoskeleton, particularly microfilaments and microtubules, plays a central role in TNT formation. Treatments that destabilize these cytoskeletal polymers markedly inhibit the formation and growth of TNT. Accordingly, cytoskeleton-destabilizing treatments also impair the heterotypic mitochondria transfer to astrocytes and the apoptotic rescue of astrocytes in the presence of pericytes. These results not only actively implicate the integrity of the cytoskeleton for the efficient intercellular exchanges through TNT but also causally link the processes of TNT formation, intercellular connections, mitochondria transport, and rescue from apoptosis to cytoskeleton integrity and dynamics aimed at preserving the BBB function.

The role of TNT in the preservation of astrocyte survival and, more generally, BBB functionality was entirely unknown. It is known that Wingless and Int-1 (WNT)/β-catenin pathway signaling regulates the BBB development,^53^ and WNT signals play a role in the endothelial cell-oligodendrocyte crosstalk.^54^ Interestingly, the WNT pathway is involved in TNT-mediated interneuronal communication.^55^ Whether WNT pathway controls the biogenesis of TNT between pericytes and astrocytes will be a topic for future investigations.

In conclusion, TNT-mediated intercellular communication between BBB cells and the TNT-mediated mitochondria transfer between healthy pericytes to post-ischemic apoptotic astrocytes could represent a new mechanism to maintain astrocyte survival and BBB functionality. This mechanism could contribute to a better understanding of the restorative role and stem cell-like behavior of pericytes during BBB recovery after ischemic stroke. Further studies will be needed to clarify the exact pathways involved in the regulation of this mechanism, to design new strategies for boosting BBB integrity, or transiently bypassing BBB for drug treatment of neurological diseases.

## Acknowledgements

We thank Dr. Elisabetta Colombo (Center for Synaptic Neuroscience, Istituto Italiano di Tecnologia, Genova, Italy) for help in confocal imaging; Drs. Diego Moruzzo and Arta Mehilli (Istituto Italiano di Tecnologia, Genova, Italy) for providing the reagents for the experiments; Rossana Ciancio and Ilaria Dallorto (Istituto Italiano di Tecnologia, Genova, Italy) for administrative help.

## Author Contributions

F.P. and V.C. designed the experiments performed co-cultures, assembloids, time-lapse and confocal microscopy, immunofluorescence, OGD-treatments, STS-treatment, apoptosis analysis, TNT, and mitochondria transfer quantification; L.S. contributed to performing OGD-treatments, TNT-quantification and perform statistical analysis. F.L. perfomed flow cytometry analysis; F.P. and V.C. wrote the manuscript; F.B. and M.S. contributed to experimental planning, data interpretation, manuscript writing and funded the research; all of the authors contributed to the final version of the manuscript.

## MATERIALS AND METHODS

### Chemicals, immunochemicals, and cell lines

Human cerebral microvascular endothelial cells (hCMEC/D3) were purchased from Cedarlane (cat. #CLU512) and cultured in endothelial basal medium (EBM-2; Lonza, cat. #190860) supplemented with 10% (v/v) fetal bovine serum (Life Technologies, cat. #10270-106), chemically defined lipid concentrate (Life Technologies, cat. #11905031), ascorbic acid (Sigma, cat. #A4544), human basic fibroblast growth factor (bFGF) (Sigma, cat. #F0291), HEPES 1M (Life Technologies, cat. #15630-080), hydrocortisone (Sigma, cat. #H0135), penicillin, 10000 units - streptomycin, 10000 μg/mL (Life Technologies, cat. #15140-122). Supplements were added according to the manual instructions. Normal human brain astrocytes (NHA) were purchased from Lonza, Clonetics (cat. #CC-2565) and cultured in ABM Astrocyte Basal Medium supplemented with SingleQuots supplements (Lonza # CC-3186) according to manual instructions. Human brain microvascular pericytes (hBVP) were purchased from ScienCell (cat. #1200) and cultured in 500 ml of PM pericyte medium supplemented with 10 ml of fetal bovine serum, 5 ml of pericyte growth supplement, and 5 ml of penicillin/streptomycin solution (Science Cell, Cat. #1201). The BBB working medium was prepared using EGM-2 Endothelial Cell Growth Medium-2 BulletKit (Lonza, cat. #CC-3162), which includes EBM-2 Endothelial basal medium and endothelial growth supplements hEGF, hydrocortisone, GA-100, hFGF-B, R3-IGF-1, ascorbic acid, heparin, and 2% (v/v) of human serum (Sigma-Aldrich, cat. #H4522). BBB working medium was used for co-culture and assembloids. In the experiments, NHA cells were used between passages 2 and 5, hBVP cells between passages 2 and 6, and hCMEC/D3 cells between passages 25 and 30. At least three independent original batches were used for each cell line to perform experiments. NHA, hBVP, and hCMEC/D3 were analyzed by immunofluorescence for the expression of the cell-specific markers GFAP, NG2, and occludin for astrocytes, pericytes, and endothelial cells, respectively (**Suppl. Figure 1**).

Staurosporine (STS, cat. #S4400), vincristine (VCR, cat. #V8879), cytochalasin D (CytoD, cat#C2618), and 4 KDa dextran-FITC (cat. #46944) were purchased from Merck. Human transferrin-CF488A conjugate was purchased from Biotium (cat. #00081). Mouse monoclonal anti-GFAP (cat. #ab10062) and rabbit polyclonal anti-NG2 (cat. #ab129051) antibodies were purchased from Abcam. The rabbit polyclonal antibody to VE-cadherin (cat. #36-1900) was purchased from Thermo Fisher.

### Live immunofluorescence analysis of 2D cell cultures

For live-labeling mitochondria, cells were incubated with 100 nM MitoTracker Deep Red FM (MT; Invitrogen, cat. #M22426) diluted in complete growth media. After 20 min of incubation, cells were extensively washed with phosphate-buffered saline (PBS), washed five times with growth medium, left overnight in the growth medium, washed with PBS, trypsinized, and centrifuged. The cell pellet was washed twice to clear the excess dye completely. After this treatment, cells were counted and co-cultured, as reported in the text. Plasma membranes were stained by exposing cells to CellMask Orange (CM; Thermo Fisher, cat. #C10045) diluted 1:1000 in FluoroBrite DMEM (Gibco, cat. #A1896701) for 10 min in the incubator, followed by extensive washes with growth medium. F-actin was stained by exposing fixed and permeabilized cells to AlexaFluor488-Phalloidin (1:500; Thermo Fisher, cat. #A12379) for 60 min at room temperature. Nuclear staining was obtained using bisbenzimide H (Hoechst 33342; Sigma cat. #B2261) at the final concentration of 10 μg/mL. To reduce the breakdown of TNT and maintain the fluorescence of MitoTracker and CellMask, cells were fixed directly in the medium by adding paraformaldehyde to a final concentration of 3.7% v/v for 15 min at 37 °C. Afterward, samples were carefully washed with PBS, permeabilized by 0.3% Triton X-100 in PBS for 15 min, blocked with 3% BSA in PBS, and incubated with primary antibody overnight at 4 °C. After careful washing, the samples were incubated with secondary antibody or fluorescent phalloidin as described above, washed in PBS, and finally mounted in Vectashield antifade mounting medium (Vector Laboratories cat. #H-1000-10).

### Cell culture treatments

#### Staurosporine

Astrocytes were treated with staurosporin (STS) at the final concentration of 1 μM for 3h in DMEM serum-free medium, washed refreshed in STS-free BBB working medium for coculture with pericytes, in the absence or presence of either cytochalasin D (CytoD, 200 nM) or vincristine (VCR, 10 nM) for 24 h.

#### Oxygen-glucose deprivation

The day before the treatment, glucose- and serum-free DMEM (DMEM no glucose, Gibco cat. #11966025) was conditioned in a hypoxic gas mixture (see below) for 24 h (OGD medium), and astrocytes were seeded on uncoated glass. The day after seeding, astrocytes were washed with OGD medium, placed in a hypoxic chamber conditioned with 2% O_2_, 5% CO_2_, 93% N_2_ (10 L/min for 5 min) in OGD medium, and incubated at 37 °C for 24 h. Thereafter, cells were gently washed and co-cultured with a constant number of pericytes in the presence or absence of CytoD (200 nM) or VCR (10 nM) for 24 h.

### Time-lapse and confocal fluorescence microscopy

For live-cell imaging, cells were seeded on confocal dishes (Thermo Fisher, cat. #150682 or Greiner Bio-One, cat. #627871), and images were acquired by Leica DMI 6000 B inverted microscope equipped for live microscopy with incubator LSM Black, temperature controller Pecon 37-2 digital, CO_2_ controller Pecon, heating unit, CTR7000 controller, Hamamatsu A3472-07 camera and light source EL6000 with a mercury metal halide bulb. To correctly visualize the mitochondria’ dynamics, images were acquired with a low-intensity light source to reduce the phototoxicity and dye photobleaching. Time-lapse images were analyzed using FIJI software

Confocal microscopy was performed using a Leica TCS SP8 or SP5 microscope. All acquisitions were performed by serial acquisition mode between frames. XYZ-series were acquired with a raster size of 1024 × 1024 in the X–Y planes and a Z-step of 0.15 μm between optical slices. Threedimensional (3D) images and projections from z-stack were constructed and processed using Leica Application Suite X software (LASX). Images were analyzed using LASX and FIJI software.

The quantification of heterotypic TNT between astrocytes and pericytes was performed as already reported.^21^ Only F-actin positive structures, detached from support and connecting astrocytes to pericytes on single confocal planes were counted. Randomly chosen images per biological replicate were acquired, and the TNT density was expressed as TNT number per field composed of 50 cells. Mitochondrial transfer in pericytes-endothelial co-cultures was analyzed on XYZ-series of confocal images using the LASX software. Pericytes mitochondria, identified by Mitotracked Deep Red staining, were counted in X-Y intracellular astrocytes planes.

### Analysis of astrocyte apoptosis

hBVP cells were cultured in 6-well plates until 50% of confluence and, the day before the analysis were treated with VCR (10 nM), CytoD (200 nM), or left untreated (controls) and stained with MT as described above. NHA (2×10^4^) were cultured in confocal dishes (Greiner Bio-One, cat. #627871) with the glass bottom without coating. NHA cells were stained with CM as described above and then either exposed to OGD for 24 h or treated with STS (1 μM for 3h). After the treatment, NHA were washed and cultured in the presence or absence of hBVP. For co-culture, hBVP were detached, and 2×10^4^ cells were seeded on top of NHA cells in each well and cultured in BBB working medium. After 24 h of co-culture, the percentage of apoptotic astrocytes was evaluated using CellEvent Caspase-3/7 Green Detection Reagent (1:250; Thermo Fisher, cat. #C10423) added directly to the medium, without washing of cells. Images were acquired using the inverted microscope used for live imaging. Random areas were acquired per biological replicate, and apoptotic astrocytes were counted.

### Flow cytometry of cell necrosis

For flow cytometry analysis, the same protocol described before for STS was applied except for the staining with MitoTracker DR and CellMask Orange. After 24h of co-culture, cells were detached (cells in suspension were also collected before the trypsinization), collected by centrifugation, resuspended in 5% FBS in PBS, counted, and propidium iodide (1 mg/ml; Thermo Fisher, cat. #P1304MP) was added directly to the medium to the final concentration of 1 μg/ml. Cells were incubated at room temperature for 10 min, protected from light and immediately analysed by flow cytometry. Measurements were then performed on MACSQuant Analyzer 10 (Miltenyi Biotec) and data analyzed with FlowJo 10.6 (Becton Dickinson & Company).

### BBB multicellular assembloids

The 3D multicellular assembloid BBB model was established as previously described.^32^ For assembloids, NHA cells were used at passage P2, hBVP cells at passages P2-P3, and hCMEC/D3 cells at passage P28. A 96-well plate was pre-coated with 50 μl of 1% agarose in PBS per well. Cells were detached, and 1.5 × 10^3^ cells per type were counted. Cell were plated in the ratio 1:1:1 in BBB working medium (200 μl total volume) in each well. After 48 h, the multicellular assembloids were self-assembled as single spheroids/well in more than 90% of the wells.

For confocal imaging, all perfect spherical assembloids were carefully collected by pipette tips (P200) in 0.2 ml PCR tube for analysis. Immunofluorescence was performed as follows. Assembloids were collected (four/tube), harvested by gravity, and immediately fixed with 3.7% (v/v) paraformaldehyde in BBB working medium for 45 min at 37 °C. Fixed assembloids were washed with PBS, permeabilized with 0.5% TritonX100 in PBS overnight at 4 °C, washed and saturated by 3% w/v bovine serum albumin (BSA) in PBS overnight at 4 °C and incubated with primary antibodies in 3% BSA (w/v in PBS) overnight at 4 °C. After incubation, assembloids were washed in 3% BSA (w/v in PBS) overnight at 4 °C, incubated with secondary antibodies for 2 h, washed three times with PBS, and deposited in glass-bottom dishes (Grenier Bio-One, cat. #627871) for confocal analysis.

For barrier integrity assays, spherical assembloids were collected in a PCR tube and immediately incubated in BBB working medium with 10 μM 4 kDa dextran-FITC (FD4; Merck, cat. #46944) or 10 μM fluorescently labeled human transferrin (Tf488; Biotium, cat. #00081) for 4 h at 37 °C under gentle agitation. The assembloids were then washed and transferred in glass-bottom dishes for confocal analysis. The permeability to FD4 and Tf488 was assessed by capturing Z-stack images, starting from the outermost surface of the assembloid towards the core. A 20x objective was used, and the fluorescence of the compounds inside the assembloid at a 50-μm depth was assumed as a proxy of assembloid permeability, as showed in a previous work.^32^

### Statistical analysis

Data are presented as means ± SE and plotting individual data points. The normal distribution of experimental data was assessed using the D’Agostino-Pearson’s normality test. When comparing two groups with normal distribution, the unpaired Student’s *t*-test was used. When more than two groups with normal distribution were compared, one-way or two-way ANOVA followed by the Tukey’s multiple comparison test were used. When more than two groups with non normal distribution were compared, Kruskal-Wallis test followed by Dunn’s multiple comparison test was used.

No predictive statistics was used to predetermine sample sizes, and the adopted sample (indicated in figure legends) were similar to those previously reported in the literature. All statistical procedures were performed using GraphPad Prism 6 software (GraphPad Software, Inc).

