## Supplemental Data 1 for "TUNNELING NANOTUBES CONNECT BLOOD-BRAIN BARRIER CELLS: ROLE OF PERICYTES IN BARRIER PRESERVATION DURING ISCHEMIA"

### SUPPLEMENTARY INFORMATION

### (Supplementary Figures)

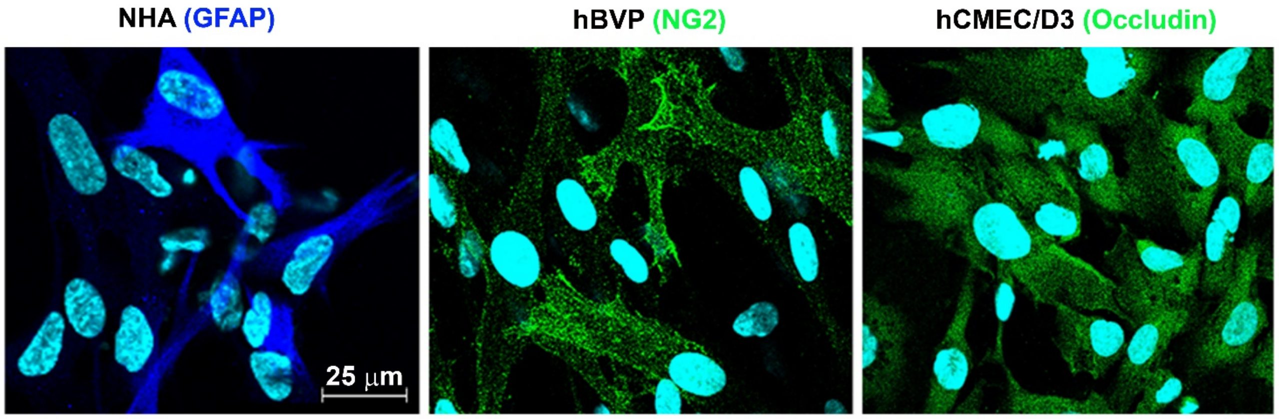


**Supplementary Figure 1. Characterization of astrocytes, pericytes and endothelial cells by specific markers.** Immunofluorescence images show the expression of GFAP in normal human astrocytes (NHA), NG2 in human brain microvascular pericytes (hBVP), and occludin in human brain microvascular endothelial cells (hCMEC/D3).

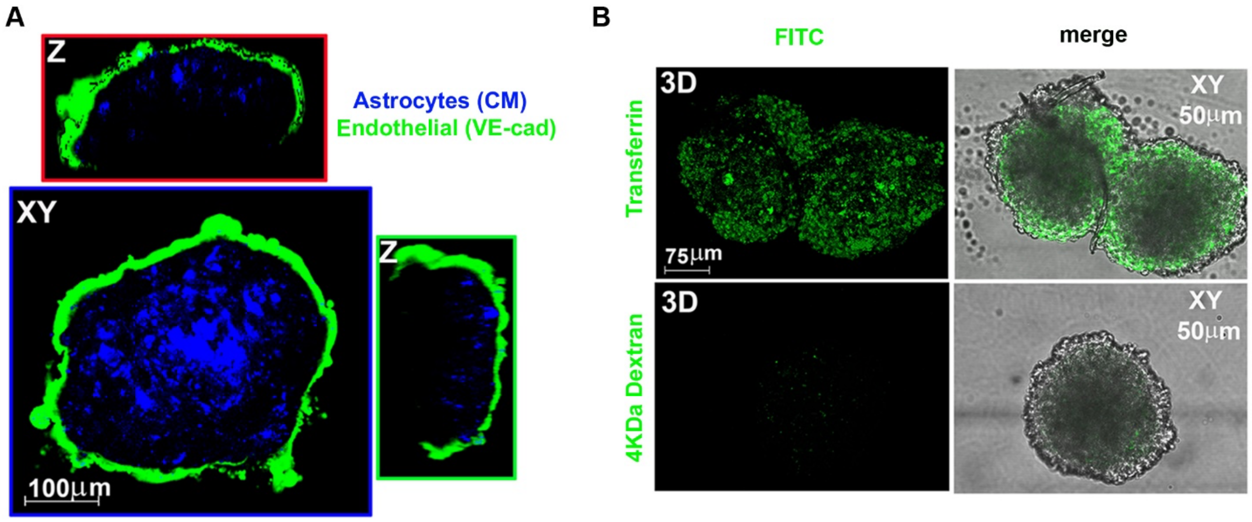


**Supplementary Figure 2. Characterization of BBB assembloids.**

**A.** Expression of VE-cadherin in BBB assembloids. Whole assembloids were analyzed by anti-VE cadherin immunofluorescence. Endothelial VE-cadherin (green) is expressed at the surface of the assembloid. CM-labeled astrocytes (blue) are detected in the core region. Representative images from N = 2 independent experiments with ≥ 12 assembloids per experiment analyzed. **B.** Barrier integrity assay in BBB assembloids. Live assembloids were incubated with either FITC-labeled 4 kDa-dextran or AlexaFluor488-labeled transferrin and analyzed by confocal microscopy[32]. Representative 3D reconstructions and Z-stack confocal fluorescence images of BBB assembloids (at a 50 µm depth) show the accumulation of transferrin (known to be BBB-permeable) within the assembloid core, while 4 kDa-dextran is excluded. Representative images from N = 4 independent experiments in which N = 6 assembloids per experiment were analyzed.

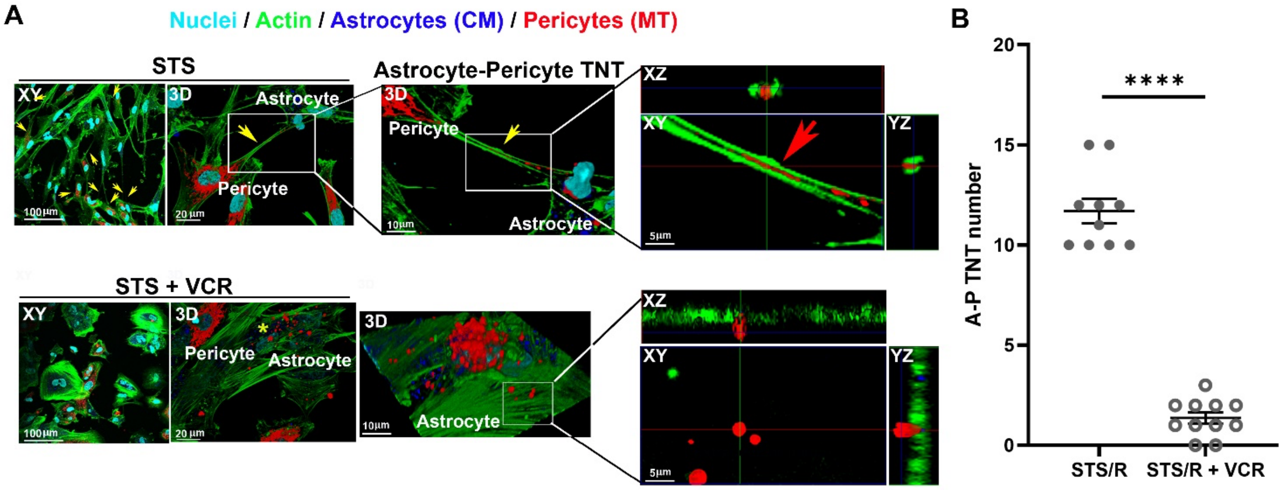


**Supplementary Figure 3. Effects of microtubule depolymerization on the STS-induced TNT formation between astrocytes and pericytes.**

**A.** CM-labeled astrocytes (blue) were treated with STS (1 µM for 3 h), washed, and co-cultured with pericyte that had been previously stained with MT (red) in the presence or absence of the the microtubule-depolymerizing drug vincristine (VCR; 10 nM). After 24 h, cells were fixed, stained for the F-actin-network with phallodin (green), and analyzed by confocal microscopy for the TNT number and mitochondria transfer. VCR prevents STS-induced astrocyte-pericyte TNT formation. Yellow arrows indicate TNT between pericytes and STS-astrocytes. The magnified insets show TNT in 3D reconstruction, XY plane and Z-projections. The red arrow indicates mitochondria inside TNT between a pericyte and an STS-astrocyte. **B.** Quantification of heterotypic-TNT (A-P) number after 24 h of co-culture. The VCR treatment strongly prevented pericyte-to-astrocyte TNT formation. N = 4 independent experiments. ****p<0.0001, unpaired Student's *t*-test.

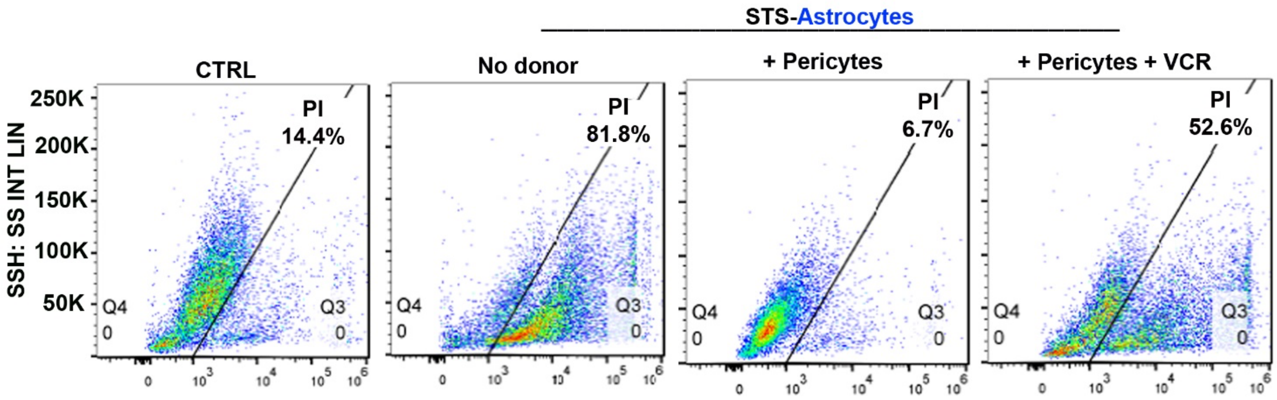


**Supplementary Figure 4. Pericytes rescue astrocytes from STS-induced necrosis.**

Astrocytes were treated with STS (1 μM for 3 h), washed, and co-cultured with pericyte in the presence or absence of VCR. After 24 h, cells were collected (cells in suspension were also collected before the trypsinization), added with propidium iodide (PI) (final concentration of 1 μg/ml) and analyzed for necrosis by flow cytometry. The percentage of PI-positive cells is shown. Pericytes strongly rescue astrocytes from STS-induced necrosis, and VCR counteracts the rescue. At least 10.000 events per condition were analyzed.
